# Quinolinic acid metabolism may mitigate AKI to CKD transition

**DOI:** 10.1101/2025.10.10.681643

**Authors:** Marie Christelle Saade, Afaf Saliba, Amanda J. Clark, Subrata Debnath, Shiqi Zhan, Nagarjunachary Ragi, Valerie Etzrodt, Rahil Al-Humaidi, Kyle Vu, Yu Tao, Esmeralda Trevino, Alejandra L Lorenzen, Guanshi Zhang, Anders H. Berg, Jason C. O’Connor, Samir M. Parikh, Kumar Sharma

## Abstract

The transition from acute kidney injury (AKI) to chronic kidney disease (CKD) remains a significant clinical problem with unclear underlying mechanisms. Emerging evidence suggests that alterations in tryptophan metabolism, particularly in the production of downstream metabolites such as quinolinic acid (QA), play a role in renal pathophysiology. QA is a NAD biosynthesis intermediate metabolized by the enzyme quinolinate phosphoribosyltransferase (QPRT). In this study, we investigated the role of QA in the AKI-to-CKD transition using experimental mouse models and clinical observations and leveraging multiple omics approaches. Systematic metabolomic profiling identified endogenous QA as one of the most significantly elevated metabolites following folic acid-(FA) induced injury. Exogenous QA exacerbated FA-induced kidney dysfunction. Conversely, aged mice deficient in QPRT showed worsened expression of kidney fibrosis markers even in absence of kidney injury, while younger littermates exhibited worsened induced kidney injury. Mice lacking QA-producing enzymes resisted experimental AKI and AKI-to-CKD progression. Multimodal spatial metabolomics analysis of human AKI kidney biopsies revealed QA accumulation in regions of inflammatory infiltration. Finally, children with CKD exhibited higher urinary QA levels compared to healthy controls. These findings underscore QA as a potential mediator of kidney injury and a therapeutic target for preventing the progression from AKI to CKD.

**One Sentence Summary:** Quinolinic acid promotes kidney damage and fibrosis, suggesting it as a contributor of AKI-to-CKD progression and a potential therapeutic target.

## INTRODUCTION

Acute kidney injury (AKI) is the sudden loss of kidney function. Even when AKI resolves, this syndrome poses significant risk for progression to chronic kidney disease (CKD). Moreover, therapeutic options are lacking, and there is an incomplete understanding of the molecular mechanisms driving this transition. (*1*) While numerous mechanisms contribute to this transition, recent work has highlighted the role of disrupted metabolic pathways, particularly those involving nicotinamide adenine dinucleotide (NAD ) biosynthesis and tryptophan metabolism, in modulating kidney injury and repair. (*2–4*) This process, known as the *de novo* NAD+ biosynthetic pathway, is the major route of enzymatic tryptophan degradation. It passes through an intermediate, quinolinic acid (QA), which constitutes the first fully committed step towards NAD+ production, whereas more upstream metabolites have multiple metabolic fates (**Fig. 1A**). QA has long been recognized for its neurotoxic effects in neurological disorders, but its impact on peripheral organs such as the kidney, remains poorly defined. (*5*) We recently identified QA as a potential contributor to renal injury in AKI and CKD (*6*) and demonstrated that overexpression of the quinolinate phosphoribosyl transferase enzyme (QPRT), which catabolizes QA, protects against cisplatin and folic acid-induced AKI. (*7*)

**Figure 1.**
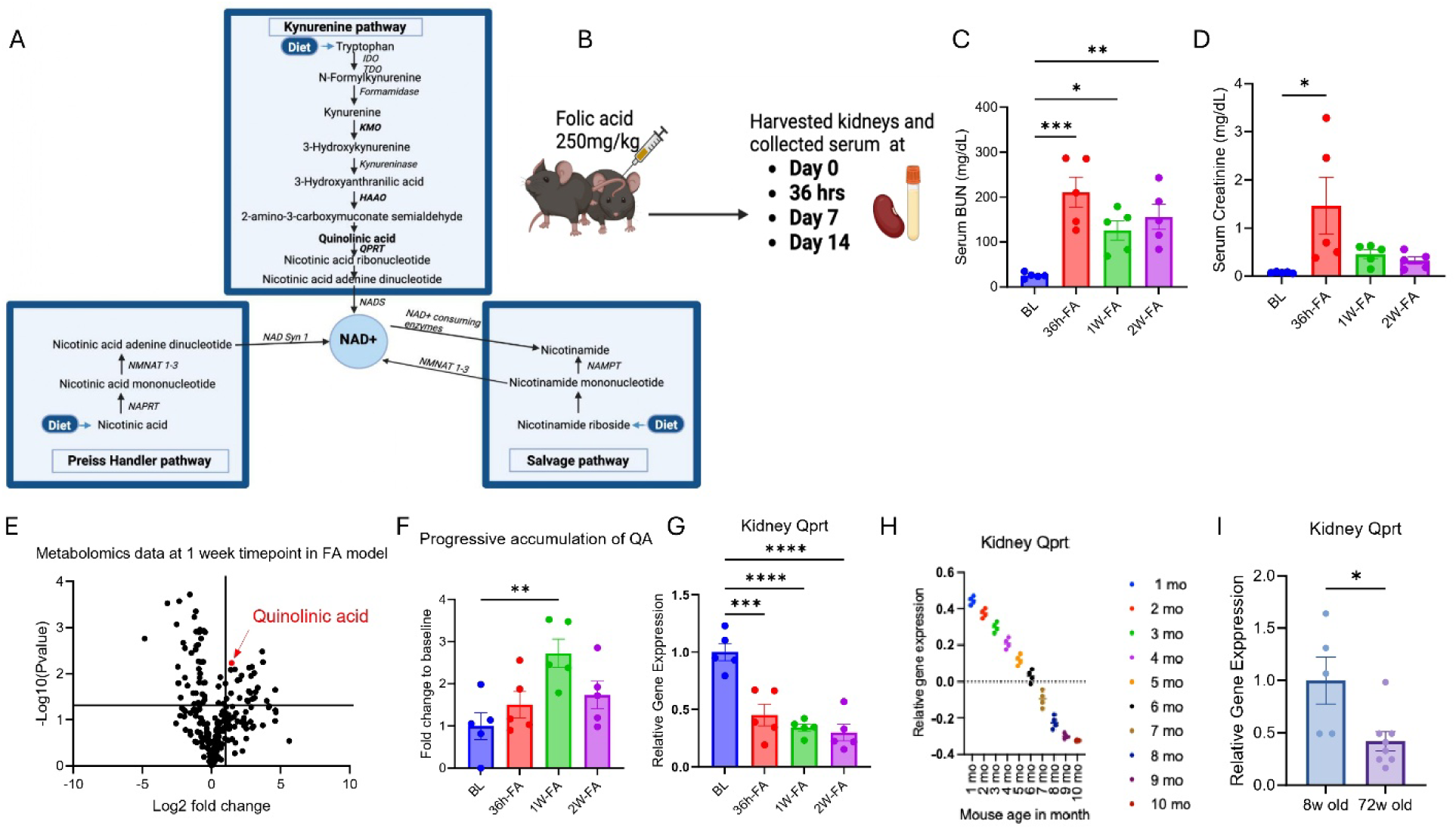
Quinolinic acid (QA) is markedly increased during AKI-to-CKD Progression, while *Qprt* expression declines with disease and aging. (**A**) Schematic overview of the NAD+ biosynthesis pathways. (**B**) Schematic representation of folic acid (FA) experiment in mice. Wild-type (WT) mice (n=5 per group) were administered FA (250 mg/kg, intraperitoneally) and sacrificed at different time points: 0, 3h, 7 days, and 14 days post injection to assess AKI-to-CKD transition. (**C**) Serum BUN (**D**) Serum creatinine. (**E**) Volcano plot showing untargeted metabolomics data at 1-week timepoint. (F) Time-course analysis of (**F**) plasma QA and (**G**) kidney *Qprt* mRNA expression. All data are presented as mean ± SEM. Statistical analysis was performed using one-way ANOVA with post hoc testing. p < 0.05 (*), p < 0.01 (**), p< 0.001 (***), p < 0.0001 (**). (**H**) Tabula Muris dataset analysis reveals that *Qprt* expression declines with age in mouse kidneys. (**I**) Kidney *Qprt* mRNA expression in 8-week-old (n=5) vs 72-week-old mice (n=8), data are presented as mean ± SEM using Mann Whitney two-tailed t-test. p < 0.05 (*).

Hence, we hypothesized that QA is not merely a metabolic intermediate in the biosynthesis of NAD+, but also a pathogenic metabolite that directly contributes to kidney injury and promotes the transition from AKI to CKD. Identifying key molecular drivers of the transition from AKI to CKD progression is crucial for developing therapeutic interventions that could halt or reverse AKI-induced damage before it evolves into CKD. Using a combination of genetic and dietary approaches in murine models, we investigated the effect of QA accumulation on this transition. Our findings were supported by human data, which further implicated QA as a pathogenic metabolite in kidney injury. Together, these results identify QA as a contributor to renal disease progression and suggest that targeting its production or promoting its clearance may offer novel therapeutic strategies to prevent the progression from acute to chronic kidney injury.

## RESULTS

### QPRT downregulation drives quinolinic acid accumulation after kidney injury

Alterations in NAD biosynthesis play a critical role in both kidney injury and aging. (*8*) Unlike in AKI models, where NAD supplementation has been shown to be beneficial, rodent models of CKD have yielded mixed results. (*9*) Based on our data, nicotinamide (NAM) supplementation in the folic acid (FA)-induced AKI-to-CKD transition model **(Fig. S1)**, did not improve outcomes. (**Fig. S2**). Therefore, we investigated other factors that might be interfering with this effect. QPRT, a key rate-limiting enzyme in this pathway, regulates the metabolism of QA and its dysregulation may contribute to pathological QA accumulation. (*10, 11*) We hypothesized that QA accumulates after kidney injury due to reduced *Qprt* expression.

To investigate this hypothesis, mice were administered with FA and sacrificed at day 0, 36 hours, 1 week, and 2 weeks post-injection for kidney and serum collection (**Fig. 1B**). Renal function was assessed via serum creatinine and BUN levels to validate the injury model. Serum BUN and creatinine were significantly elevated at 36 hours, 1 week, and 2 weeks after FA administration, compared to baseline levels, confirming the successful induction of kidney injury (**Fig. 1C and 1D**). We performed systematic metabolomics analysis of kidney tissues, which revealed that QA was among the most significantly upregulated metabolites, with levels peaking at the one-week time point after FA administration (**Fig. 1E and 1F**). Kidney qPCR analysis showed a marked downregulation of *Qprt* expression following injury across all three post-injury time points (**Fig. 1G**) indicating a metabolic bottleneck likely resulting from QPRT downregulation.

### QPRT expression decreases with age

Given that aging is a strong and poorly understood risk factor for AKI, (*12*) we investigated *Qprt* expression with age. An analysis of the Tabula Muris dataset, a public database, shows a decline of *Qprt* with age in kidney tissue (**Fig. 1H**). We validated those findings in our lab by comparing kidneys from 8-week-old and 72-week-old mice, which showed that *Qprt* expression decreases with age (**Fig. 1I**). Together, these results demonstrate that *Qprt* expression is significantly reduced both in kidney injury and with aging, contributing to the accumulation of QA, a potentially toxic intermediate in the NAD biosynthesis pathway.

### Endogenous QA accumulation exacerbates the AKI-to-CKD transition in mice and impairs kidney health in chronic settings

In previous work, we demonstrated that genetic reduction of *Qprt*, and potentially QA accumulation, exacerbates the severity of ischemic and nephrotoxic-AKI in mice. (*3, 7*) Building on this, we tested whether sustained QA accumulation due to impaired QPRT activity may drive the AKI-to-CKD transition. We assessed kidney injury in *Qprt ^+/-^* mice compared to *Qprt ^+/+^* littermates 14 days after FA injection. (**Fig. 2A**). *Qprt*^+/-^ mice exhibited significantly higher BUN (**Fig. 2B**) and creatinine levels (**Fig. 2C**), along with increased mRNA levels of *Col3a1* (**Fig. 2D**) and *Tgfb* (**Fig. 2E**).

**Figure 2.**
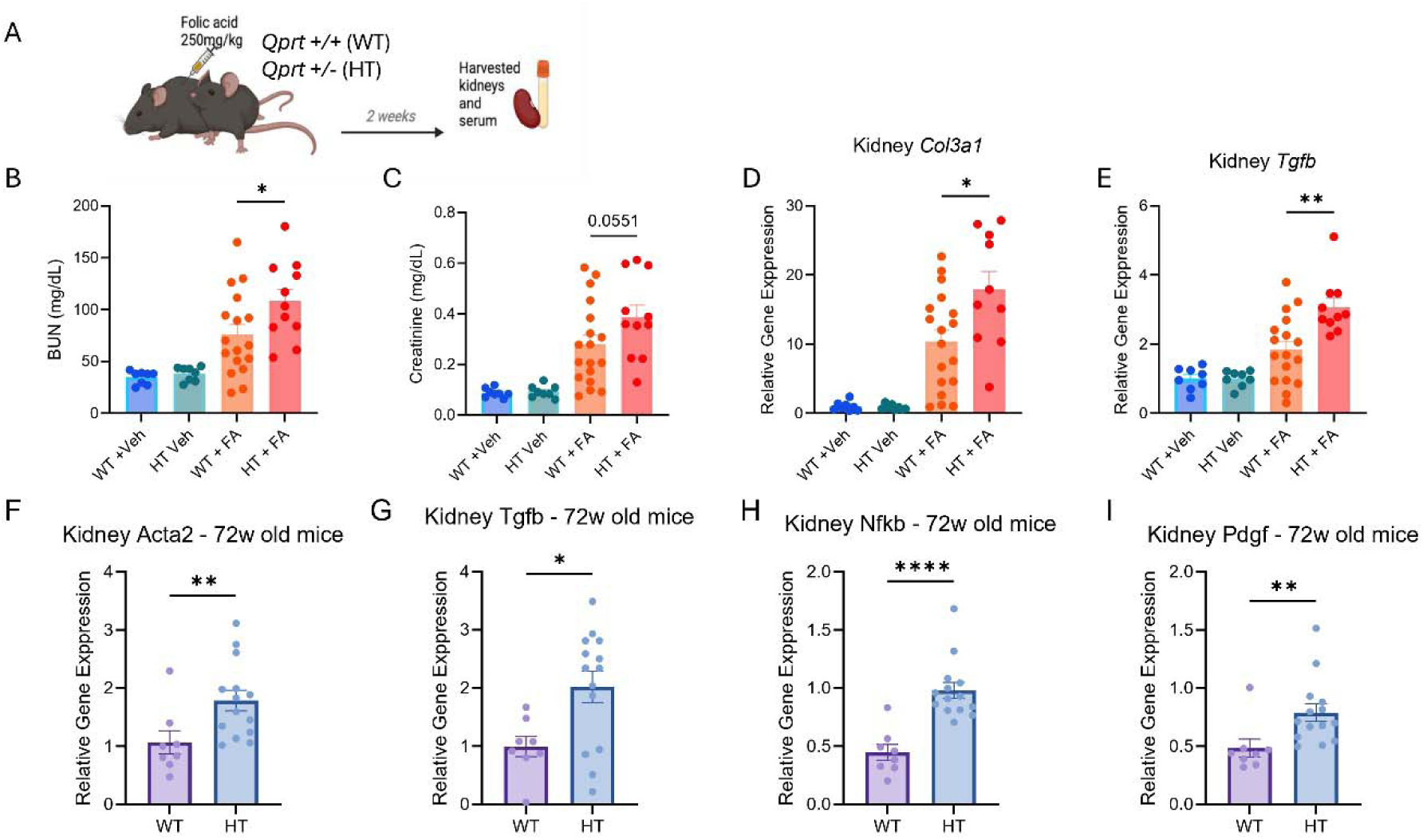
Endogenous quinolinic acid accumulation exacerbates the AKI-to-CKD transition in mice and impairs kidney health in chronic settings. (**A**) Schematic representation of a folic acid (FA)-induced AKI-to-CKD transition experiment: *Qprt ^+/-^*(HT) mice (n=11) and *Qprt* ^+/+^ (WT) (n=18) were injected with FA (250mg/kg) on day 0 and were sacrificed 14 days later. (**B**) Serum BUN (**C**) Serum creatinine (**D**) Kidney collagen, type III, alpha 1 (*Col3a1)* mRNA expression (**E**) transforming growth factor beta (*Tgfb)* mRNA expression. Analysis of pro-fibrotic and pro-inflammatory markers in kidneys of 72-week-old *Qprt ^+/-^* (HT) mice and *Qprt* ^+/+^ (WT) including (**F**) Actin, alpha 2 (*Acta2)* mRNA expression (**G**) *Tgfb* mRNA expression (**H**) Nuclear Factor kappa-light-chain-enhancer of activated B cells (*Nfkb)* mRNA expression (**I**) Platelet-Derived Growth Factor (*Pdgf)* mRNA expression in kidneys. All data are presented as mean ± SEM. Statistical analysis was performed using Mann Whitney two-tailed t-test. p < 0.05 (*), p < 0.01 (**), p< 0.001 (***), p < 0.0001 (****).

Furthermore, since aging is a physiological model of chronic kidney dysfunction and is associated with declining *Qprt* expression (**Fig. 1**), we investigated whether QPRT insufficiency also accelerated pathological changes associated with kidney aging. We examined 72-week-old *Qprt* ^D/D^ and *Qprt* ^D/D^ kidneys for expression of pro-fibrotic and pro-inflammatory markers in the absence of injury. 72-week-old *Qprt* ^D/D^ demonstrated no difference in serum BUN or creatinine (**Fig. S3**) but had a marked upregulation of fibrosis and inflammation-associated genes compared to *Qprt* ^D/D^ littermates. Specifically, expressions of *Acta2*, *Tgfb*, *Nfkb*, and *Pdgf* were significantly higher (**Fig. 2F-I**), reinforcing the role of QPRT in maintaining kidney health during aging by clearing QA accumulation.

### Exogenous QA accumulation exacerbates the AKI-to-CKD transition in mice

To directly assess the pathogenic potential of QA, we next administered exogenous QA to mice and evaluated its impact on kidney function and AKI-to-CKD progression. QA was provided for 14 days in drinking water at a concentration of 0.5 g/L (**Fig. 3A**), inducing excess plasma QA (**Fig. 3B**) at levels that are comparable to those we previously reported in CKD patients. (*6*) Mice that were administered with QA exhibited significantly elevated serum creatinine levels compared to control mice (**Fig. 3C**).

**Figure 3.**
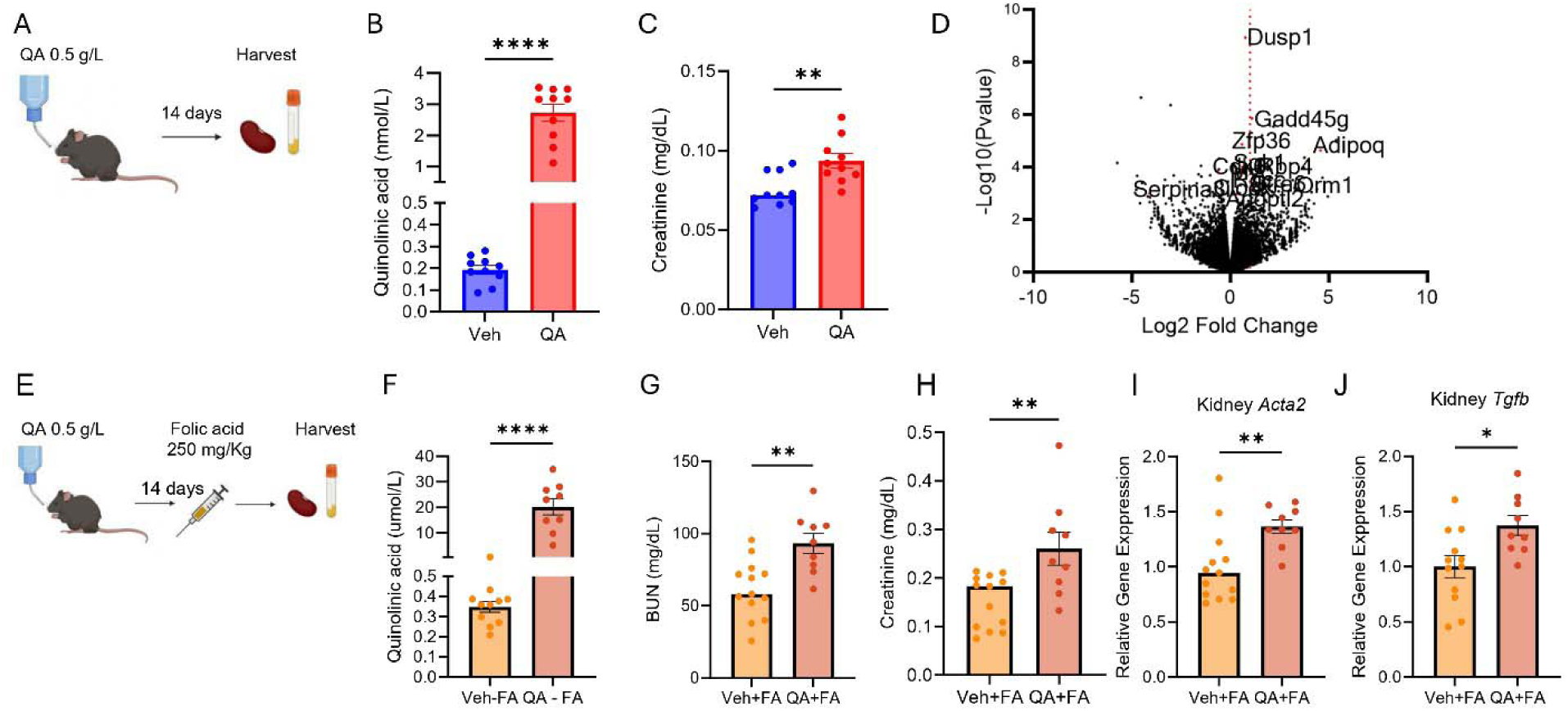
Exogenous quinolinic acid (QA) accumulation impairs kidney health and exacerbates the AKI-to-CKD transition in mice. (**A**) Schematic representation of QA administration through drinking water in a mouse experiment: WT mice (n=10 per group) received QA 0.5g/L in their drinking water vs. regular water, harvested 14 days later. (**B**) plasma QA (**C**) Serum creatinine (**D**) Kidney RNA sequencing analysis. (**E**) Schematic representation of QA administration through drinking water in a FA CKD mice experiment: WT mice (n=9-13 per group) received QA 0.5g/L in their drinking water vs regular water for 14 days, injected with FA (250mg/kg, intraperitoneally) at day 14, harvested at day 28. (**F**) plasma QA levels (**G**) Serum BUN and (**H**) creatinine (**J**) kidney *Acta2* and (**K**) *Tgfb* mRNA levels. All data are presented as mean ± SEM. Statistical analysis was performed using Mann-Whitney 2 tailed t-test. p < 0.05 (*), p < 0.01 (**), p< 0.001 (***), p < 0.0001 (****).

To further investigate the molecular effects of QA, we performed RNA sequencing on kidney tissues from five representative mice per group. A volcano plot of differentially expressed genes revealed that 14 of the top 100 genes (ranked by p-value) are implicated in kidney disease. (**Fig. 3D**). The full list includes Dusp1 (Dual specificity phosphatase 1), Gadd45g (Growth arrest and DNA-damage-inducible protein GADD45 gamma), Sgk1 (Serum/glucocorticoid regulated kinase 1), Rbp4 (Retinol binding protein 4), Myc (MYC proto-oncogene, bHLH transcription factor), Rgcc (Regulator of cell cycle), Orm1 (Orosomucoid 1), Stra6 (Stimulated by retinoic acid 6), Bmal1 (Brain and muscle ARNT-like 1), Angptl2 (Angiopoietin-like 2), Adipoq (Adiponectin, C1Q and collagen domain-containing), Cdk6 (Cyclin-dependent kinase 6), Serpina3i (Serine protease inhibitor A3I), and Clock (Circadian locomotor output cycles kaput) are involved in various biological processes. Among these, Gadd45g, Rbp4, Orm1, Stra6, and Adipoq were upregulated by more than 2-fold (**Table S1**). Collectively, these expression changes support a pathological environment marked by metabolic dysregulation, oxidative stress, and maladaptive repair in the kidney.

We separately evaluated the effect of chronic exogenous QA exposure in mice subjected to FA. In this experiment, wild type mice received QA (0.5g/L) in drinking water for 14 days, followed by IP injection of FA on day 14 and sacrificed on day 28. (**Fig. 3E**). Mice that received both FA injection and QA supplementation showed significantly higher levels of serum QA, creatinine, and BUN, as well as increased kidney mRNA levels of *Acta2* and *Tgfb1*, compared to mice that only received FA (**Fig. 3F-J**).

### Reduction of QA improves renal outcomes in a mouse model of AKI-to-CKD

To investigate the contribution of QA to maladaptive kidney repair, we administered FA to knockout mice that lack key enzymes upstream of QA production. Specifically, 3-hydroxyanthranilate 3,4-dioxygenase (HAAO) and kynurenine 3-monooxygenase (KMO). HAAO catalyzes the conversion of 3-hydroxyanthranilic acid to QA, representing the final step in the pathway leading to QA production. KMO, acts earlier in the pathway by converting kynurenine to 3-hydroxykynurenine, thereby channeling metabolites toward QA synthesis (**Fig. 1A**). At baseline, wild type (WT), *Haao*^-^/^-^, and *Kmo*^-^/^-^ mice, displayed normal body weights, BUN and creatinine levels indicating absence of an overt baseline phenotype. As expected, *Kmo*^-^/^-^ mice exhibited accumulation of kynurenine (in plasma and kidney) blockade at the KMO step and both *Haao*^-^/^-^, and *Kmo*^-^/^-^ mice had significantly lower QA levels confirming pathways blockade (**Fig. S4**). When administered with FA for 14 days (**Fig. 4A**) WT mice exhibited significant increase in plasma and kidney QA levels while *Haao^-/-^* and *Kmo^-/-^* mice exhibited significantly reduced QA levels (**Fig. 4B-C**). Suppression of QA accumulation was associated with marked improvements in BUN levels and plasma creatinine (**Fig. 4D-E**) further evidence that QA may contribute to CKD progression.

**Figure 4.**
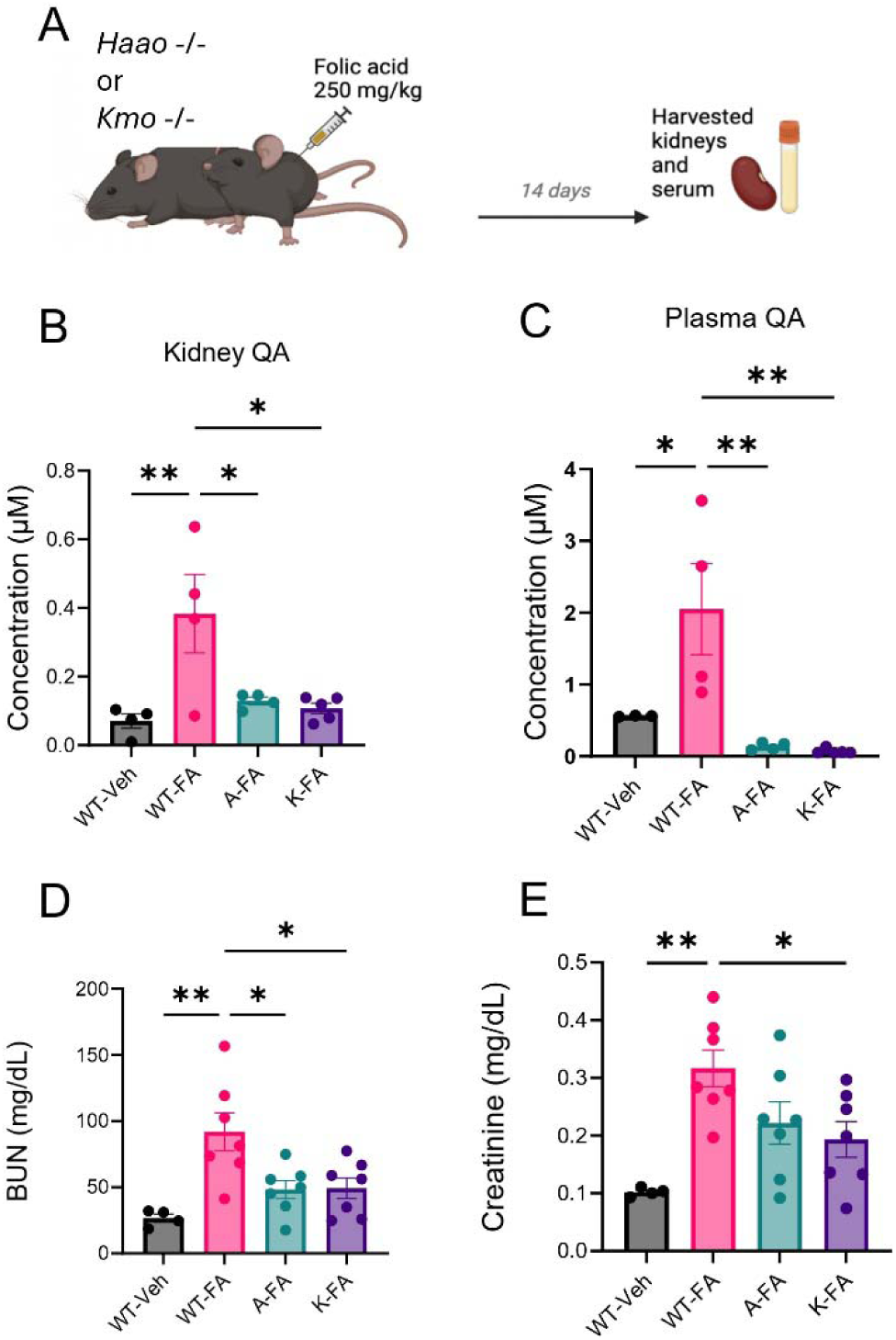
Quinolinic acid (QA) reduction improves kidney function in a mouse model of AKI-to-CKD Transition. (**A**) Schematic of the folic acid (FA)-induced AKI-to-CKD transition model. Wild-type (WT, *Haao*^-^/^-^ (denoted “A”), and Kmo^-^/^-^ (denoted “K”) mice were administered FA (250 mg/kg, intraperitoneally), and kidneys were collected 14 days post-injection (n = 4-7 per group). (**B-C**) QA levels in kidney and plasma, measured by LC-MS. (**D-E**) BUN and serum creatinine levels. Data are presented as mean ± SEM. Statistical significance was determined using one-way ANOVA followed by appropriate post hoc tests. p < 0.05 (*), p < 0.01 (**), p < 0.001 (***), p < 0.0001 (****).

### Mice deficient in QA are protected from cisplatin-induced AKI

We also sought to test whether *Haao*^-^/^-^, and *Kmo*^-^/^-^ mice would also have resistance to nephrotoxic-induced AKI but here we administered a single intraperitoneal dose of cisplatin to separate groups of knockouts and WT mice (25 mg/kg) and sacrificed 72 hours later to assess kidney injury (**Fig. 5A**). QA levels were significantly increased in cisplatin-administered WT mice and significantly reduced in both kidney tissue and plasma of cisplatin-administered *Haao*^-^/^-^ and *Kmo^-/-^*mice (**Fig. 5B-C**). Suppression of QA synthesis was associated with marked renal protection. Cisplatin administered *Haao*^-^/^-^ and *Kmo^-/-^*mice exhibited significantly lower BUN and serum creatinine levels, compared to Cisplatin administered WT mice (**Fig. 5D-E**), which was associated with significantly reduced mRNA levels of the kidney injury marker lipocalin 2 (*Lcn2*) in both knockout strains (**Fig. 5F**). These results show that QA ablation is partially protective from experimental nephrotoxic-AKI.

**Figure 5.**
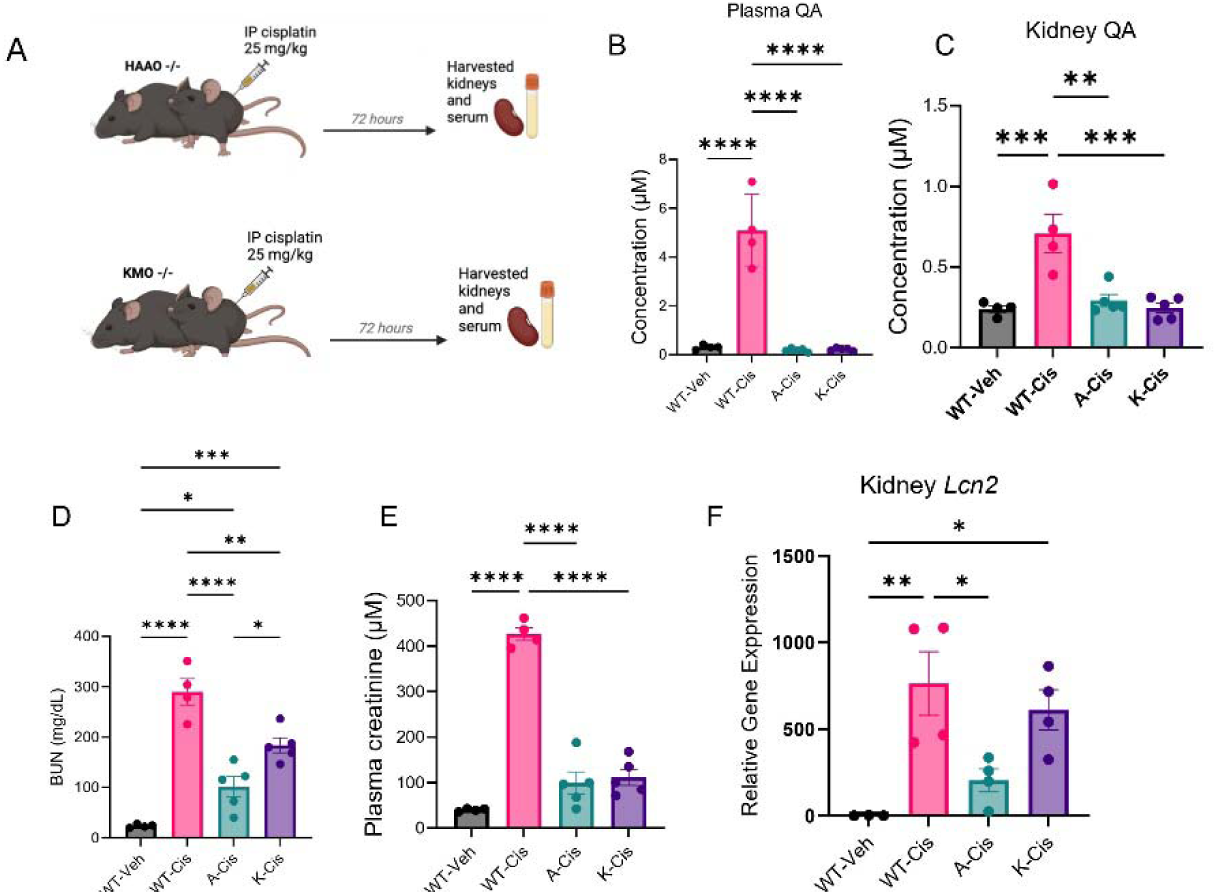
Mice lacking quinolinic acid (QA) producing enzymes exhibit resistance to cisplatin-induced AKI. (**A**) Schematic representation of the experimental design: wild-type (WT), *Haao*^-^/^-^ (dented “A”), and Kmo^-^/^-^ (denoted “K”) mice (n = 4-5) were administered cisplatin (25 mg/kg, intraperitoneally) and sacrificed 72 hours. **(B-C)** QA levels in plasma and kidney tissues. (**D)** BUN and (**E)** serum creatinine levels. (**F**) Lipocalin-2 (*Lcn2*) mRNA expression in kidney cortex samples. All data are presented as mean ± SEM. Statistical analysis was performed using one-way ANOVA with post hoc testing. p < 0.05 (*), p < 0.01 (**), p< 0.001 (***), p < 0.0001 (****).

### QA accumulates in injured human kidneys

To assess the translational relevance of our findings, we analyzed human kidney biopsies from patients with AKI and healthy reference tissue (HRT) using matrix-assisted laser desorption/ionization mass spectrometry imaging (MALDI-MSI) multi-modal analysis. Biopsies were obtained from the Kidney Precision Medicine Project (KPMP) and included 8 HRT samples and 19 AKI samples (**Table S2)**. Histological assessment by periodic acid-Schiff (PAS) staining demonstrated preserved architecture in HRT and characteristic tubular injury and cellular infiltration in AKI kidneys (**Fig. 6A-B**). Ion images revealed differential spatial distribution of tryptophan pathway metabolites. While tryptophan and kynurenine were not significantly increased (**Fig. 6C-H**), QA exhibited marked accumulation specifically within injured areas in AKI samples (**Fig. 6I-K**). Notably, QA localized predominantly to regions of inflammatory infiltration, as shown by co-registered overlays of QA ion signals with serial PAS-stained sections (**Fig. 6L-Q**).

**Figure 6.**
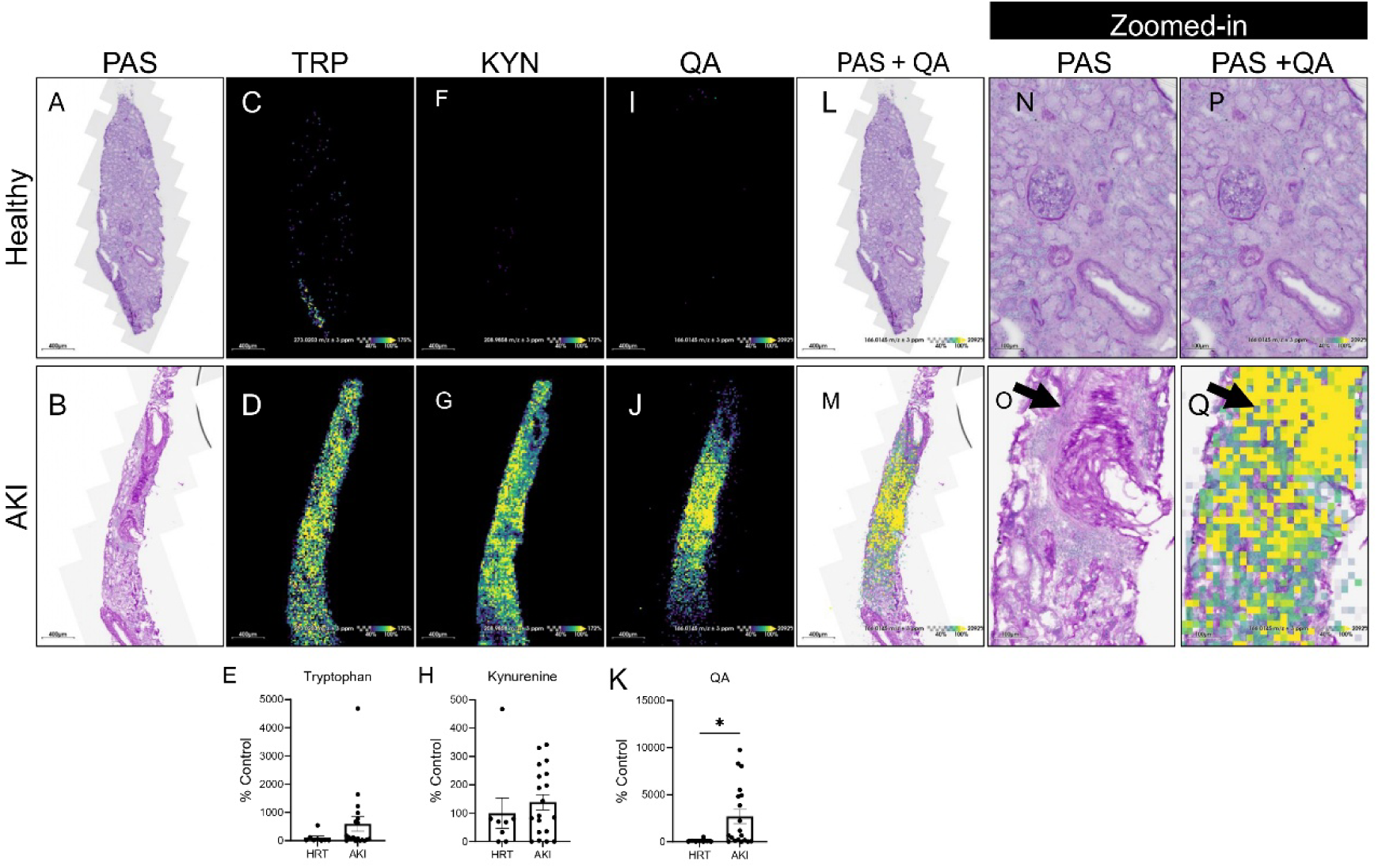
Quinolinic acid (QA) accumulates in regions of inflammatory infiltration in human AKI kidney tissue. Biopsies were obtained from healthy reference (HRT) (n = 8) and acute kidney injury (AKI) (n = 19) patients via the Kidney Precision Medicine Project (KPMP). MALDI-MSI was performed at the Center for Precision Medicine. **(A-B)** PAS-stained kidney sections **(C-K)** Ion images of tryptophan (Trp), kynurenine (KYN), and quinolinic acid (QA) and their corresponding intensity quantification presented as percentage (%) of control (HRT). Data are shown as mean ± SEM. Statistical comparisons were made using unpaired two-tailed t-tests. p < 0.05 (*). **(L-M)** Co-localization of PAS and QA with optical image overlaid on ion signal of serial sections. **(N-O)** Zoomed-in views of (A-B) **(P-Q)** Zoomed-in views of (L-M). Healthy reference kidney tissues (top) and AKI kidney tissues (bottom).

### QA is increased in children with CKD and is highly dialyzable in patients with ESKD

Next, we sought to evaluate urinary QA levels in children with CKD compared to those without known kidney disease, who were undergoing urine testing for routine screening or unrelated clinical indications. We included 72 pediatric patients with CKD and 36 controls (**Table S3**). To account for variations in urine concentration, QA levels were normalized to urinary tryptophan. The urinary QA to tryptophan ratio was significantly higher in patients with CKD compared to healthy controls. (**Fig. 7A**) Urine protein-to-creatinine ratio (UPCR) was significantly elevated in patients with CKD confirming the presence of impaired kidney function (**Fig. 7B**).

**Figure 7.**
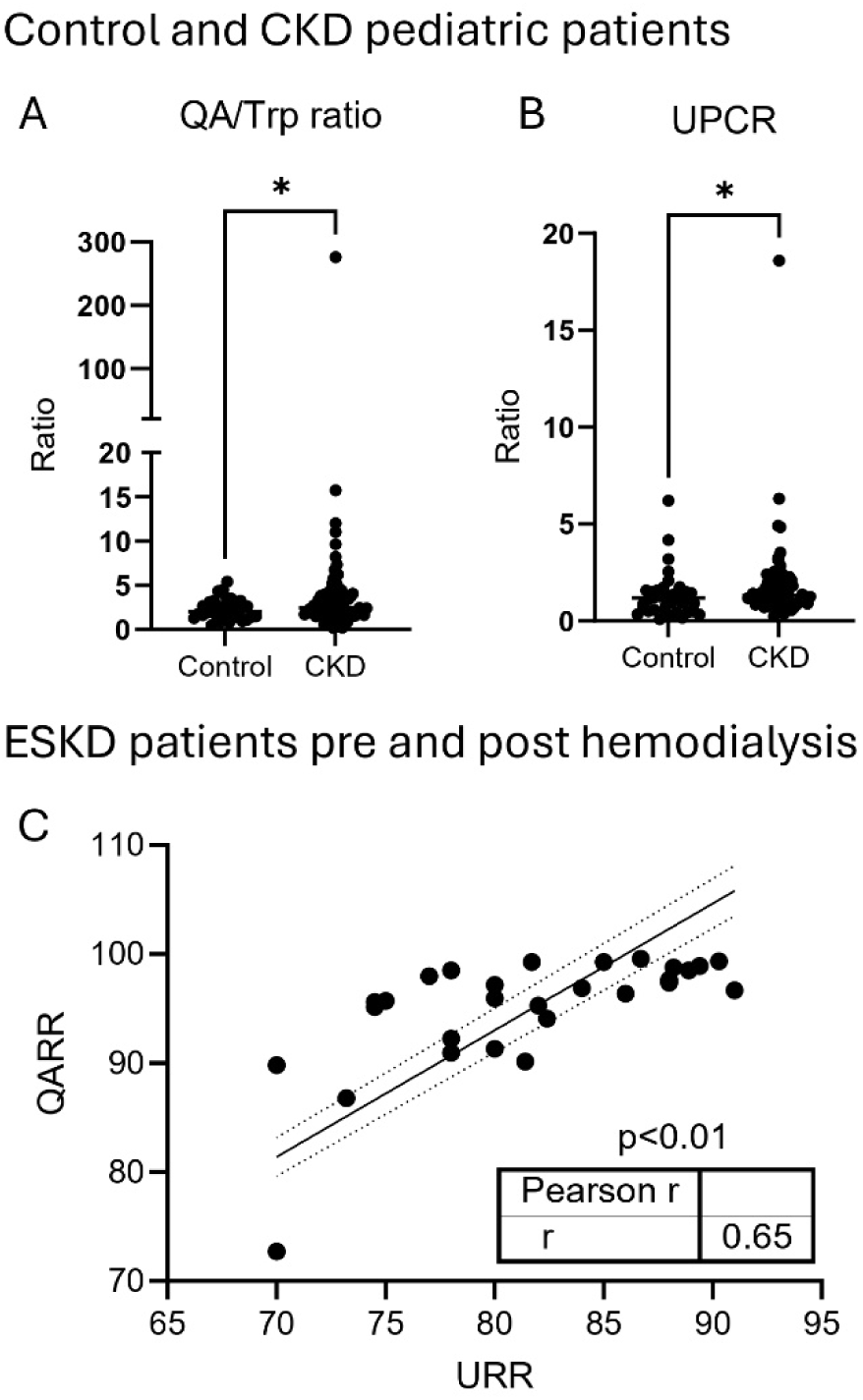
Quinolinic acid (QA) in chronic kidney disease (CKD) and end stage kidney disease (ESKD) (**A**) QA to tryptophan ratio (QA/Trp) in control in CKD vs. CKD participants. (**B**) Urinary Protein to Creatinine ratio (UPCR) in CKD vs. control participants. 72 pediatric participants with CKD and 36 control participants. Data presented as mean ± SEM. Statistical analysis was performed using Mann Whitney two-tailed t-test. p < 0.05 (*) (**C**) Pearson correlation between plasma QA reduction ratio (QARR) and urea nitrogen reduction ratio (URR) before and after a single hemodialysis session in 29 adult participants with end stage kidney disease.

Lastly, to evaluate the dialyzability of QA and related tryptophan metabolites, we measured plasma levels before and after a single hemodialysis session in 29 ESKD patients. As shown in **Tables S4 and S5**, QA exhibited a high reduction ratio (QARR: 94.84% ± 5.44), closely mirroring that of blood urea nitrogen (URR: 81.35% ± 6.06), indicating that QA is effectively cleared during dialysis. QA reduction ratio (QARR) correlated significantly with URR (**Fig. 7C**), indicating that QA behaves similarly to dialyzable small molecules.

## DISCUSSION

In this study, we identify QA as a pathogenic metabolite that contributes to AKI and its progression to CKD. It is one of the most significantly upregulated metabolites identified through an unbiased approach in the AKI-to-CKD transition model. Its accumulation in *Qprt^+/-^* mice, where QPRT is the enzyme responsible for catalyzing QA, exacerbated kidney injury in AKI-to-CKD progression models, supporting a role for QA as a nephrotoxic metabolite. This was further demonstrated with exogenous administration of QA to WT mice, at concentrations comparable to those observed in the serum of humans with CKD, induced renal dysfunction, as mice had transcriptional activation of inflammatory and oxidative stress pathways. Genetic deletion of QA-producing enzymes (HAAO and KMO) protected mice from cisplatin-induced AKI and FA-induced AKI-to-CKD transition. QA accumulation is evident in human subjects across the spectrum of kidney injury including AKI, CKD, and ESKD. Multimodal mass spectrometry imaging analysis of human kidney biopsies with AKI showed unique accumulation of QA at regions of inflammatory cells, suggesting it may be associated with inflammation. In CKD, we found that urinary QA also accumulates in a pediatric cohort, providing the first evidence of its elevation in children, a population generally characterized by fewer comorbidities than adults. Notably, QA was also found to be efficiently removed by a single session of hemodialysis, further supporting its relevance as a modifiable uremic toxin.

One potential mechanism by which QA promotes maladaptive repair is through activation of fibrogenic pathways. Our data show that elevated QA levels or decreased *Qprt* levels are associated with increased expression of fibrosis-related genes in injured or aged kidneys. This finding aligns with our previous work demonstrating that pathophysiological concentrations of QA upregulate fibrotic markers in human kidney cortical cells (HK-2) cells. (*6*)

QA itself may be directly nephrotoxic, even in the presence of NAD precursors. In fact, prior studies have described QA-induced neurotoxicity via glutamate receptor overactivation and ROS generation. (*13–15*) Furthermore, although QA can be converted to NAD , this reaction is limited by the expression and enzymatic capacity of QPRT, which may become saturated under pathologic conditions. Thus, both decreasing QA production and enhancing its clearance may be necessary for therapeutic benefit. Studies targeting upstream enzymes in the kynurenine pathway, such as KMO and IDO, are already underway in inflammatory, neurological and oncologic contexts. (*16, 17*) Similarly, we and others showed that NAD repletion strategies, including supplementation with nicotinamide riboside or nicotinamide mononucleotide confer protection in AKI and aging. (*3, 9, 18–20*)

We acknowledge limitations of our study, particularly the use of germline, whole-body knockout models. Future experiments with tissue-specific inducible effects will further support the renal relevance of this pathway. In summary, we identify QA as a potential driver of AKI-to-CKD transition, as both endogenous and exogenous accumulation of this metabolite lead to worsened kidney function. Our findings also suggest that the accumulation of QA exerts a protective role in this transition. Nevertheless, additional studies are warranted to confirm and validate these outcomes in different models and in human populations.

## MATERIALS AND METHODS

### Study design

This study combined controlled laboratory experiments with observational analyses of human clinical samples. The experimental arm used murine models to investigate the role of QA in the transition from AKI-to-CKD, demonstrating that QA accumulation is nephrotoxic and that its clearance can mitigate disease progression. The observational arm evaluated QA accumulation and clearance in human kidney disease, incorporating urinary QA analysis in pediatric CKD patients and healthy controls, multimodal histological assessment of kidney biopsies from AKI patients and healthy reference tissue (HRT), and plasma metabolite profiling in dialysis patients to assess QA dialyzability.

Sample sizes were based on prior data, known variability, and literature benchmarks, with power analyses performed where applicable (power = 0.8, α = 0.05). Each condition included a minimum of 4 biological replicates (individual mice).

In experimental mouse models studies only, males are employed. In human studies, samples from both males and females were employed; however, no direct comparison between the sexes was made due to the limited sample size.

### Animal studies

All experiments described in Figures 1-4 were conducted in compliance with the NIH Guide for the Care and Use of Laboratory Animals and approved by IACUC from the University of Texas Southwestern Medical Center. All experiments described in Figures 5 and 6 were carried out after IACUC approval from the University of Texas Health Science Center at San Antonio.

### FA-induced injury in wild type (WT) mice on a C57BL/6J background (8-weeks old) (**Fig. 1**)

Male mice were injected IP with FA (#216630100) 250 mg/kg dissolved in 0.3 M sodium bicarbonate, euthanized at different time point (day 0, 36h, 7 days, 14 days) post-injection, plasma and kidneys were collected. WT mice on C57B6J background we sacrificed at 8-weeks vs. 72-weeks old for kidney tissue analysis.

***Qprt* ^+/-^ mice (Fig. 2**) were previously described (C57BL/6J background). (*3*) Male mice (8-12 weeks old) were injected intraperitoneally (IP) with FA (#216630100) 250 mg/kg dissolved in 0.3 M sodium bicarbonate. Mice were euthanized 14 days post-injection, and plasma and kidneys were harvested for further analysis.

Male C57BL/6J WT mice (72-week-old) were sacrificed for analysis, plasma and kidneys were collected. (**Fig. 3**), mice were given QA in drinking water (0.5 g/L) for 14 days, with water exchanged every other day. On day 14, the mice were euthanized, and plasma and kidneys were collected.

In a separate cohort, C57BL/6J mice received QA (0.5g/L) in drinking water for 14 days, followed by IP injection of FA at day 14 and mice were sacrificed on day 28. Plasma and kidneys were collected.

***Kmo****^-^/^-^* **and *3Haao****^-^/^-^* **mice (Fig. 4 and 5**): Mouse studies were carried out after IACUC approval from the University of Texas Health Science Center at San Antonio.

Experiments utilized wild-type (WT) mice as well as KMO and HAAO transgenic mice (Kmotm1a(KOMP)Wtsi and Haaotm1a(KOMP)Wtsi) on a C57BL/6N background were designed and generated by the Mouse Biology Program (MBP, www.mousebiology.org) at the University of California Davis (UC Davis) and were provided by lab of Dr. Jason O’Connor. (*21, 22*)

Male mice (8-12 weeks old) were injected IP with FA (#216630100) 250 mg/kg dissolved in 0.3 M sodium bicarbonate, as described previously.(*23*) Mice were euthanized 14 days post-injection, and plasma and kidneys were harvested for further analysis.

Male mice (8-12 weeks old) were injected IP with 20 mg/kg cisplatin dissolved in saline solution, as described previously. (*24*) Mice were euthanized 72h later, and plasma and organs were collected.

### Biochemical Analysis

Blood urea nitrogen (BUN) levels were measured from plasma samples using a colorimetric detection kit (Arbor Assays BUN Detection Kit #K024-H or EIA-BUN kit Invitrogen), following the manufacturers’ protocol.

Serum creatinine was measured using capillary electrophoresis at the University of Texas Southwestern renal physiology core.

Serum QA was measured using commercial ELISA (Immusmol - IS-I-0100R)

### RNA and qPCR

RNA was isolated from cell or tissue lysate using Bio-Rad Aurum Total RNA Mini Kit. cDNA was made using Bio-rad iScript Advanced cDNA Synthesis Kit and Thermo Fisher Scientific RevertAid Reverse Transcription Kit (Cat# 4374966) with 1 µg of RNA for each sample reaction. qPCR master mix was prepared by mixing SYBR Green PCR Master Mix (Thermo Fisher Scientific, Cat# A25780). The primers sequences are listed in the supplementary methods. **Metabolomics**

### Untargeted metabolomics

Metabolomics were performed by Children’s Research Institute (CRI) core facility at the University of Texas Southwestern Medical Center.

### Targeted bulk metabolomics

Tryptophan metabolites were measured from plasma, frozen kidney cortex lysates (10 µL) using ZipChip (908 Devices, Boston, MA) coupled with mass spectrometry or by bulk LC/MS analysis at the Center for Precision Medicine, University of Texas Health Science Center at San Antonio. (*6*) The plasma (10 µL) samples were treated in the same manner as the tissue lysates. A microfluidic chip that integrates capillary electrophoresis (CE) with nano-electrospray ionization through a ZipChip interface separates metabolites. LC/MS analyses were performed on the Thermo Q Exactive HF-X Orbitrap mass spectrometer (Thermo, San Jose, CA) interfaced with heated electrospray ionization source (HESI) and coupled with Thermo Vanquish HPLC system. An aliquot of 2.5 µL of the sample was injected into the instrument using an autosampler. The chromatographic separation took place on an Agilent ZORBAX HILIC PLUS column with 3.5µm particle size and with the dimensions 2.1 ×100 mm with a phase composition of 10 mM ammonium formate, 0.05% formic acid in Millipore water (component A), and 0.05% formic acid in 95% acetonitrile (component B) at 0.3 mL/ min flow rate was used. (*25*)

Quinolinic acid was analyzed with the XBridge Peptide BEH C18 column with 2.5µm particle size and with the dimensions 2.1 ×100 mm with a phase composition of 10 mM ammonium acetate, and 0.05 % acetic acid in 100% acetonitrile (component B) at 0.3 mL/ min flow rate was used. Data acquisition and processing was carried out using Thermo Scientific’s Xcalibur Quant Browser software.

### RNA sequencing

RNA sequencing was performed by McDermott Center Next Generation Sequencing Core at the University of Texas Southwestern Medical Center.

### Human kidney biopsies

Fresh frozen human kidney biopsy tissues from HRT (n = 8) and AKI (n = 19) participants were procured from consented patients through the Tissue Procurement Service at the University of Michigan Ann Arbor (UMich) as part of the Kidney Precision Medicine Project (KPMP) consortium (https://kpmp.org/). Samples collected for this study received exempt approval from the UMich Institutional Review Board due to anonymization. The study adhered to all relevant ethical regulations and guidelines. Participant characteristics are included in **Table S2**.

### Matrix-assisted laser desorption/ionization mass spectrometry imaging (MALDI-MSI) of kidney biopsy

Frozen kidney biopsy specimens obtained from participants in the KPMP were cryosectioned at a thickness of 7 μm using a Leica CM1950 cryostat. Tissue sections were mounted onto indium tin oxide (ITO)-coated glass slides for MALDI-MSI and onto standard glass slides for histological analysis. A MALDI matrix solution consisting of 1,5-diaminonaphthalene (DAN; 5.6 mg/mL in 5% HCl and 50% ethanol) was applied to the ITO-mounted sections using the HTX M3+ automated sprayer under the following conditions: flow rate of 20 μL/min, 25 passes, 10 psi liquid nitrogen sheath gas, and a spraying velocity of 1200 mm/min. MALDI-MSI was conducted using a Thermo Q Exactive HF-X Orbitrap mass spectrometer integrated with a UV-laser MALDI source (Spectroglyph LLC). Data acquisition was performed in negative ion mode across an m/z range of 70–1000 with a spatial resolution of 20 μm. Raw mass spectrometry data were converted to .ibd and .imzML formats using ImageInsight software and subsequently uploaded to METASPACE, an online platform for molecular annotation of MALDI-MSI datasets. Adjacent serial sections of the kidney cortex were fixed in 4% formaldehyde post-sectioning and stained with periodic acid–Schiff (PAS) stain, processed in University of Texas Health Science Center at San Antonio. Autofluorescence and brightfield (AF-BF) images of PAS-stained sections were acquired using a Zeiss Axioscan 7 digital slide scanner equipped with a 20x objective lens.

### Quinolinic acid, tryptophan, and kynurenine measurement in human kidney cortex

The spatial distribution of quinolinic acid, tryptophan, and kynurenine in frozen kidney biopsy sections was analyzed using Bruker SCiLS Lab software. MALDI-MSI datasets for each sample were imported into SCiLS, and corresponding pre-MSI AF images were registered to the total ion current (TIC) image using a two-point alignment method. PAS–stained images from adjacent serial sections were subsequently overlaid onto the AF images. Kidney tissue regions were defined based on PAS-stained images, with structure confirmed via AF imaging. Regions of interest (ROIs) were defined to comprehend the entire cortex biopsy, and ion intensity data corresponding to quinolinic acid (m/z 166.0146 ± 3 ppm), tryptophan (m/z 273.0203 ± 3 ppm), and kynurenine (m/z 208.9858 ± 3 ppm) were extracted for each pixel within the ROI following TIC normalization. Mean ion intensity values for quinolinic acid were calculated by averaging pixel-level intensities within the ROI. To confirm molecular identity, tandem mass spectrometry (MALDI-MS/MS) was performed on frozen kidney cortex sections targeting the quinolinic acid, tryptophan, and kynurenine precursor ion. The resulting fragment ion m/z values were verified by Human Metabolome Database (HMDB) spectral library.

### Pediatric cohort of patients with CKD

This study was approved by the institutional review board at University of Texas Southwestern (STU-2021-0835) and was carried out in accordance with the Declaration of Helsinki. This study was performed on discarded samples with minimal risk to patients, thus no informed consent was obtained.

All urine samples discarded from a tertiary care children’s hospital were collected during a four-month period. Electronic medical records were reviewed, and patients who met criteria were sorted into 2 categories: 1. Patients with CKD (formally diagnosed based on the medical record) (n=72); 2. Outpatient controls who had urine samples sent for routine screening test (n=36).

Urine samples were sent to the Berg lab at Cedars Sinai Health Sciences University and QA was measured using Targeted Mass Spectrometry. Urine creatinine measured using a commercial assay (Bioassay systems, DICT-500) was used to normalize QA to mice urine concentration.

### Hemodialysis study

**Study Population.** In this cross-sectional study, we enrolled clinically stable prevalent end stage kidney disease (ESKD) patients who had been receiving thrice-weekly in-center hemodialysis for at least 3 months (Supplemental Table AS2). Patients with an acute infection requiring hospitalization within the past 30 days and documented chronic hepatitis were excluded. All patients received standard management for ESKD and associated comorbidities as per the recommended guidelines. (*26*) All patients were dialyzed for an average of 4 hours with high-flux polysulfone dialyzers using bicarbonate based dialysate. Blood flow ranged between 300 and 400 mL/min and dialysate flow at a constant rate of 800 mL/min.

The study protocol was approved by the Institutional Review Board, and written informed consent was obtained from all participants prior to any study procedures. University of Texas Health San Antonio Institutional Review Board Approval #20110429HU.

### Data Collection

Blood specimens were collected in EDTA tubes from the arterial limb of the arteriovenous fistula immediately prior to administration of heparin (pre-dialysis) and again at the end of dialysis session (post-dialysis). Collected blood was centrifuged at 2,500 rpm for 20 min. Plasma was aliquoted and stored at −80°C until analyzed. To minimize variabilities, all study procedures were performed during the mid-week hemodialysis treatment. Pre- and Post-dialysis BUN levels were quantitatively determined using the enzymatic conductivity rate method (Roche Cobas 6000, Indianapolis, IN 46250).

### HPLC

Tryptophan and selective metabolites in the kynurenine (KYN) pathway were measured in plasma by liquid chromatography/mass spectrometry (LC-MS) as reported previously. (*27*) Briefly, 50μL plasma was diluted with 5 times of 0.2% acetic acid. Stable isotope–labeled standards, 2-picolinic-d4 acid, 2,3-pyridinedicarboxylic acid-d3, L-TRP-13C11,15N2, and KYN, were added at the time of extraction as internal standards for absolute quantification. The diluted samples were vortexed and transferred to 0.5-mL Millipore Amicon Ultra filter (3 kDa). The filter tubes were centrifuged at 13 500g for 60 minutes at 4°C and the extracts were transferred to glass vials for LC-MS analyses. High-performance liquid chromatography/electrospray ionization mass spectrometry (HPLC-ESI-MS) analyses were conducted on a Thermo Fisher Q Exactive mass spectrometer with online separation by a Thermo Fisher/Dionex Ultimate 3000 HPLC. HPLC conditions were as follows: column, YMC-Pack ODS-AQ, 3 μm, 2 × 100 mm (YMC; Allentown, PA); mobile phase A, 0.5% formic acid in water; mobile phase B, 1% formic acid in acetonitrile; flow rate, 200 μL/min; gradient, 1% B to 30% B for 5 minutes and held at 70% B for 5 minutes to clean the column. The MS analyses were conducted using full MS scan (70 000 resolution) with positive ion detection. Standard curves were generated for all targeted compounds using appropriate stable isotope–labeled internal standards and native compounds. Quantitative results were obtained by reference of the experimental peak area ratios to the standard curves.

### Calculation

The reduction rate (RR) or percentage of removal of a metabolite was calculated according to the following equation:

RR (%) = (Cpre – Cpost) / Cpre X 100, where Cpre and Cpost are the plasma concentrations of a metabolite before and after hemodialysis procedure, respectively.

The urea reduction rate (URR) is a quantitative measurement of a hemodialysis patient’s urea clearance during a single dialysis session based on three parameters: the clearance or removal of urea from the blood by a dialyzer, dialysis duration or the length of a dialysis treatment or session, and the volume of distribution of urea in a patient.

### Statistical Analyses

Continuous variables were compared with two-tailed t-test or Mann-Whitney test, while multiple group comparisons were performed using one-way ANOVA test followed by Tukey test for multiple comparisons (details included in figures legends). p-value of less than 0.05 was considered statistically significant. All statistical analyses were conducted using Prism 9 software. For hemodialysis study, differences between metabolites were analyzed with t-test. A two-sided p value <0.05 was considered statistically significant with analyses performed using the SAS software (version 9.2; SAS Institute, Cary, NC, USA). Correlations were analyzed using simple linear regression.

## Acknowledgments

We thank Natalia Kuhn in the lab of Dr. Jason C. O’Connor for technical support, including provision of Kmo^-/-^ and *Haao*^-/-^ mice and assistance with genotyping. We thank Dr. Vamsidhara Vemireddy for their support and assistance.

We gratefully acknowledge the Kidney Precision Medicine Project (KPMP) for providing access to human kidney biopsy samples. MALDI-MSI analysis was performed at the Center for Precision Medicine, Kumar Sharma Lab, in accordance with approved KPMP protocols and data use agreements. We thank the KPMP consortium for their commitment to open science and for enabling this research.

We gratefully acknowledge the essential contributions of our patient participants and the support of the American public through their tax dollars.

The authors acknowledge the University of Michigan Medical School Central Biorepository (RRID:SCR_026845) for providing biospecimen storage, management, and distribution services in support of the research reported in this publication/grant application/presentation.

## KPMP grant acknowledgment

KPMP is supported by the National Institute of Diabetes and Digestive and Kidney Diseases (NIDDK) through the following grants: U01DK133081, U01DK133091, U01DK133092, U01DK133093, U01DK133095, U01DK133097, U01DK114866, U01DK114908, U01DK133090, U01DK133113, U01DK133766, U01DK133768, U01DK114907, U01DK114920, U01DK114923, U01DK114933, U24DK114886, UH3DK114926, UH3DK114861, UH3DK114915, and UH3DK114937.

The content is solely the responsibility of the authors and does not necessarily represent the official views of the National Institutes of Health

## Author contributions

Conceptualization: AS, MCS, SMP, KS

Methodology: AS, MCS, SMP, KS

Investigation: AS, MCS, AJC, SD, SZ, NR, VE, KV, RH, YT, ET, AL, GZ, AB

Resources: JOC, SMP, KS Visualization: AS, MCS

Funding acquisition: AS, SMP, KS Project administration: SMP, KS

Supervision: SMP, KS

Writing – original draft: AS, MCS

Writing – review & editing: SMP, KS, AS, MCS, AJC, SD

## Data and materials availability

All data are available in the main text or the supplementary materials. RNA seq data deposition in progress.

## References

1. S. G. Coca, S. Singanamala, C. R. Parikh, Chronic kidney disease after acute kidney injury: a systematic review and meta-analysis. Kidney Int 81, 442–448 (2012).

2. K. M. Ralto, E. P. Rhee, S. M. Parikh, NAD(+) homeostasis in renal health and disease. Nat Rev Nephrol 16, 99–111 (2020).

3. A. Poyan Mehr et al., De novo NAD(+) biosynthetic impairment in acute kidney injury in humans. Nat Med 24, 1351–1359 (2018).

4. M. T. Tran et al., PGC1alpha drives NAD biosynthesis linking oxidative metabolism to renal protection. Nature 531, 528–532 (2016).

5. R. Schwarcz, W. O. Whetsell, Jr., R. M. Mangano, Quinolinic acid: an endogenous metabolite that produces axon-sparing lesions in rat brain. Science 219, 316–318 (1983).

6. A. Saliba et al., Quinolinic acid potentially links kidney injury to brain toxicity. JCI Insight 10, (2025).

7. A. J. Clark et al., Hepatocyte nuclear factor 4alpha mediated quinolinate phosphoribosylltransferase (QPRT) expression in the kidney facilitates resilience against acute kidney injury. Kidney Int 104, 1150–1163 (2023).

8. A. J. Clark, M. C. Saade, S. M. Parikh, The Significance of NAD+ Biosynthesis Alterations in Acute Kidney Injury. Semin Nephrol 42, 151287 (2022).

9. R. Alhumaidi, H. Huang, M. C. Saade, A. J. Clark, S. M. Parikh, NAD(+) metabolism in acute kidney injury and chronic kidney disease transition. Trends Mol Med 31, 669–681 (2025).

10. H. S. Youn et al., Structural Insights into the Quaternary Catalytic Mechanism of Hexameric Human Quinolinate Phosphoribosyltransferase, a Key Enzyme in de novo NAD Biosynthesis. Sci Rep 6, 19681 (2016).

11. M. C. Saade, A. J. Clark, S. M. Parikh, States of quinolinic acid excess in urine: A systematic review of human studies. Front Nutr 9, 1070435 (2022).

12. J. C. Peng et al., Development of mortality prediction model in the elderly hospitalized AKI patients. Sci Rep 11, 15157 (2021).

13. R. Lugo-Huitron et al., Quinolinic acid: an endogenous neurotoxin with multiple targets. Oxid Med Cell Longev 2013, 104024 (2013).

14. R. G. Tavares et al., Quinolinic acid inhibits glutamate uptake into synaptic vesicles from rat brain. Neuroreport 11, 249–253 (2000).

15. W. M. Behan, M. McDonald, L. G. Darlington, T. W. Stone, Oxidative stress as a mechanism for quinolinic acid-induced hippocampal damage: protection by melatonin and deprenyl. Br J Pharmacol 128, 1754–1760 (1999).

16. M. Platten, E. A. A. Nollen, U. F. Rohrig, F. Fallarino, C. A. Opitz, Tryptophan metabolism as a common therapeutic target in cancer, neurodegeneration and beyond. Nat Rev Drug Discov 18, 379–401 (2019).

17. M. M. Krupa, T. Pienkowski, A. Tankiewicz-Kwedlo, T. Lyson, Targeting the kynurenine pathway in gliomas: Insights into pathogenesis, therapeutic targets, and clinical advances. Biochim Biophys Acta Rev Cancer 1880, 189343 (2025).

18. J. Giroud-Gerbetant et al., A reduced form of nicotinamide riboside defines a new path for NAD(+) biosynthesis and acts as an orally bioavailable NAD(+) precursor. Mol Metab 30, 192–202 (2019).

19. S. He et al., NAD(+) ameliorates endotoxin-induced acute kidney injury in a sirtuin1-dependent manner via GSK-3beta/Nrf2 signalling pathway. J Cell Mol Med 26, 1979–1993 (2022).

20. Q. Song et al., The Safety and Antiaging Effects of Nicotinamide Mononucleotide in Human Clinical Trials: an Update. Adv Nutr 14, 1416–1435 (2023).

21. J. M. Heisler, J. C. O’Connor, Indoleamine 2,3-dioxygenase-dependent neurotoxic kynurenine metabolism mediates inflammation-induced deficit in recognition memory. Brain Behav Immun 50, 115–124 (2015).

22. J. M. Parrott et al., Neurotoxic kynurenine metabolism is increased in the dorsal hippocampus and drives distinct depressive behaviors during inflammation. Transl Psychiatry 6, e918 (2016).

23. L. J. Yan, Folic acid-induced animal model of kidney disease. Animal Model Exp Med 4, 329–342 (2021).

24. S. M. Sears, A. Orwick, L. J. Siskind, Modeling Cisplatin-Induced Kidney Injury to Increase Translational Potential. Nephron 147, 13–16 (2023).

25. P. Pallerla et al., Evaluation of amino acids and other related metabolites levels in end-stage renal disease (ESRD) patients on hemodialysis by LC/MS/MS and GC/MS. Anal Bioanal Chem 415, 6491–6509 (2023).

26. F. National Kidney, KDOQI Clinical Practice Guideline for Hemodialysis Adequacy: 2015 update. Am J Kidney Dis 66, 884–930 (2015).

27. S. Debnath et al., Tryptophan Metabolism in Patients With Chronic Kidney Disease Secondary to Type 2 Diabetes: Relationship to Inflammatory Markers. Int J Tryptophan Res 10, 1178646917694600 (2017).

