## Supplementary materials for "Quinolinic acid metabolism may mitigate AKI to CKD transition"

**Supplementary Figure S1:** Folic acid (FA)-induced mouse model of AKI-to-CKD transition.

**Supplementary Figure S2:** Nicotinamide supplementation in the folic acid-induced AKI-to-CKD transition model.

**Supplementary Figure S3:** Serum creatinine and BUN levels in 72-week-old *Qprt* $^{+/+}$ (WT) vs. *Qprt* $^{+/-}$ (HT) mice.

**Supplementary Figure S4:** Renal injury and quinolinic and acid levels in *Haao- Kmo*-Knockout mice compared to wild-type.

**Supplementary Table S1:** Top differentially expressed genes in kidneys of mice administered with 0.5 g/L quinolinic acid in drinking water for 14 days.

**Supplementary Table S2:** Metadata for selected healthy reference (HRT) and acute kidney injury (AKI) samples from Kidney Precision Medicine Project (KPMP)

**Supplementary Table S3**: Pediatric chronic kidney disease (CKD) study participants characteristics

**Supplementary Table S4:** End stage kidney disease study participants characteristics

**Supplementary Table S5:** Data analysis of reduction ratios of BUN and kynurenine pathway metabolites after a single hemodialysis session in the end stage kidney disease study

**Supplementary Methods**

**Supplementary Figure S1: Folic acid (FA)-induced mouse model of AKI-to-CKD transition.** Wild type mice were injected with vehicle (n=10) or FA 250mg/kg (n=12) on day 0. Animals were sacrificed on day 14. (A) Serum creatinine, (B) BUN and (C) *Tgfb* mRNA expression in kidneys.

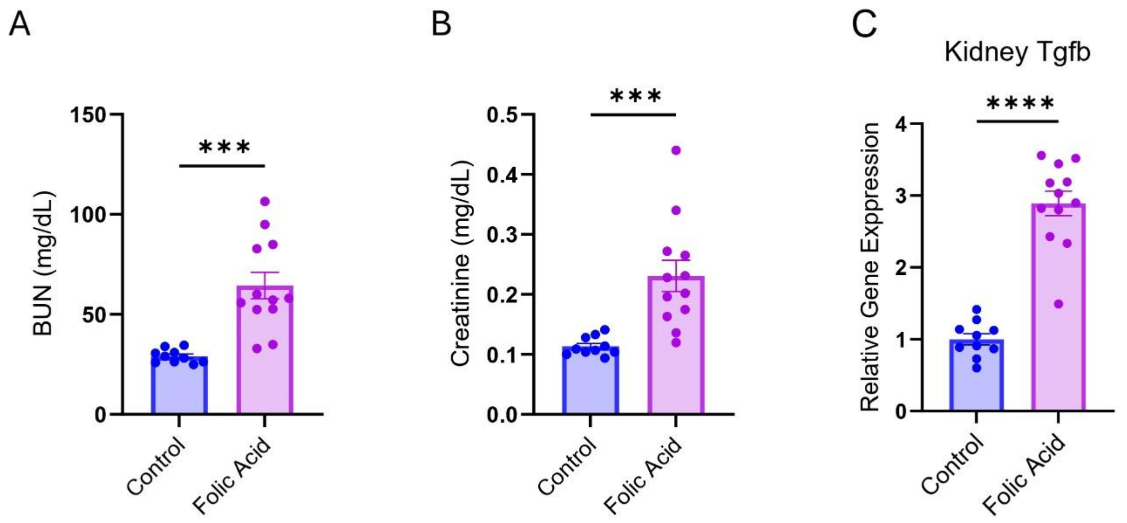

**Supplementary Figure S2: Nicotinamide supplementation in the folic acid-induced AKI-to-CKD transition model.** Mice were administered with 0.6% NAM in drinking water starting on day 0, injected intraperitoneally with 250 mg/kg FA on the same day and sacrificed on day 14 (n=5-8 per group). Metabolites were measured by untargeted metabolomics, including plasma nicotinamide (NAM) (**A**), creatinine (**B**), urate (**C**). Renal *Acta2* (**D**) and *Lcn2* (**E**) mRNA expression. All data are presented as mean ± SEM. Statistical analysis was performed using one-way ANOVA with post hoc testing. p < 0.05 (*), p < 0.01 (**), p < 0.001 (***).

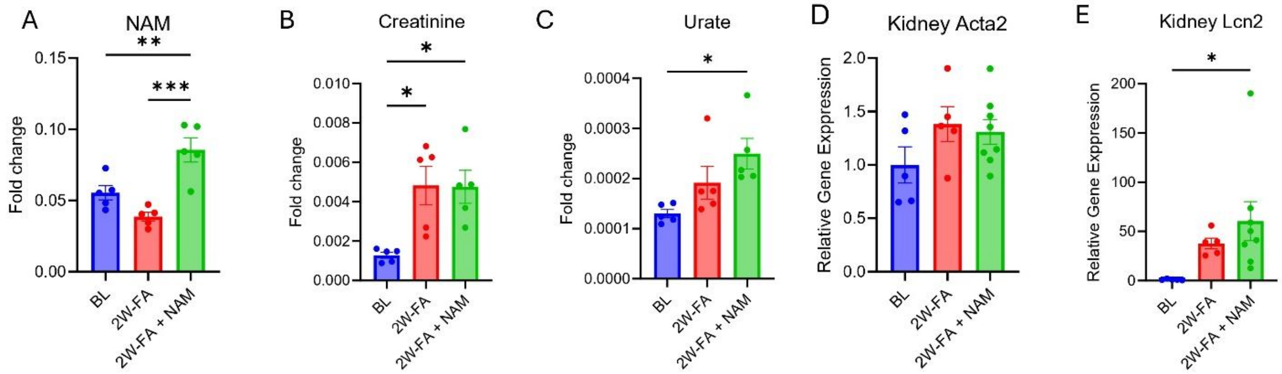

**Supplementary Figure S3: Serum creatinine and BUN levels in 72-week-old *Qprt* $^{+/+}$ (WT) vs. *Qprt* $^{+/-}$ (HT) mice. - related to Figure 3** (**A**) Serum creatinine and (**B**) BUN in aged *Qprt*$^{+/+}$ (WT) vs. *Qprt* $^{+/-}$ (HT) mice (n=7-14 per group). All data are presented as mean ± SEM.

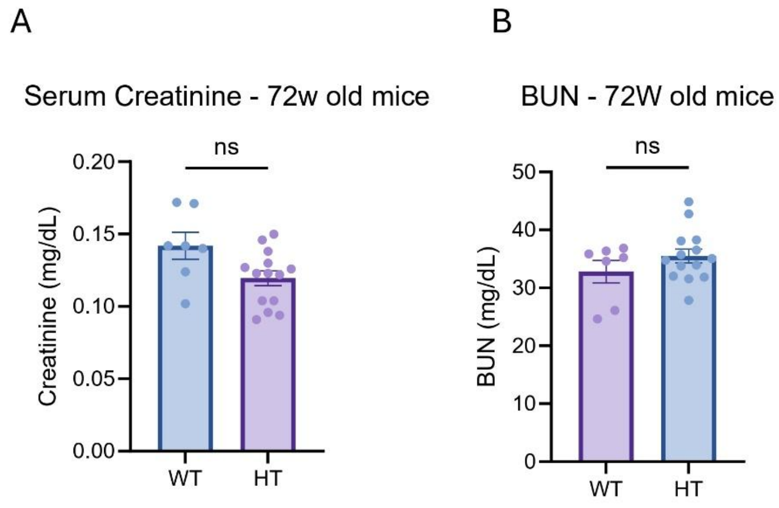

**Supplementary Figure S4: Renal injury and quinolinic and acid (QA) levels in Haao- Kmo-Knockout mice compared to wild-type. (A)** BUN levels, **(B)** Plasma creatinine levels**,** (**C**) plasma tryptophan **(D)** plasma kynurenine, (**E**) plasma QA, **(F)** kidney tryptophan **(G)** kidney kynurenine, (**H**) Kidney QA levels in wild-type (WT), *Haao* $^{-/-}$ (denoted "A") and *Kmo*$^{-/-}$ (denoted "K") n=4 per group C57BL/6N background. All data are presented as mean ± SEM. Statistical analysis was performed using one-way ANOVA. $p < 0.05$ (*), $p < 0.01$ (**), $p < 0.0001$ (****).

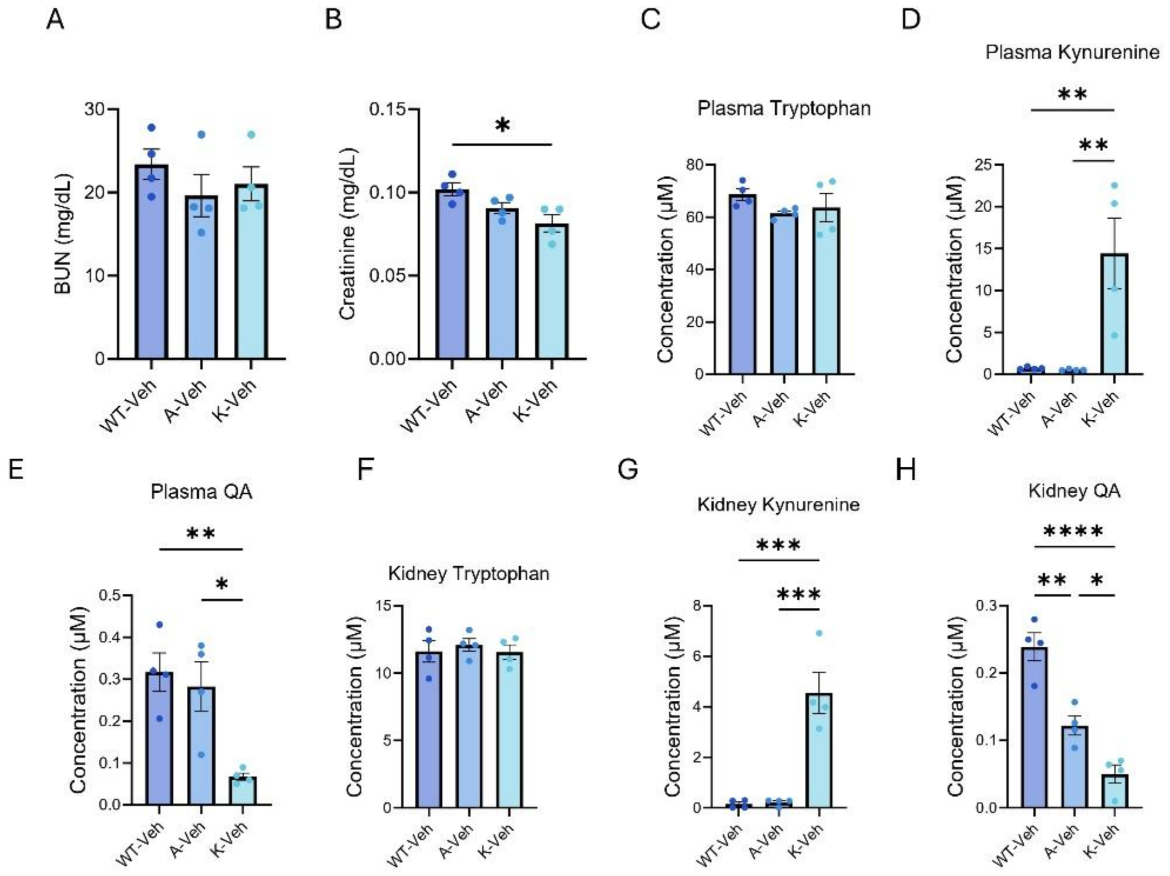

**Supplementary Table S1.** Top differentially expressed genes in kidneys of mice administered with 0.5 g/L quinolinic acid in drinking water for 14 days.

| **Gene abbreviation** | **Gene full name** | **Log2 Fold change** | **Base -10 logarithm of p-value** |
|---|---|---|---|
| Dusp1 | Dual Specificity Phosphatase 1 | 0.772965198 | 8.93287844 |
| Gadd45g | Growth Arrest and DNA Damage Inducible Gamma | 1.102903637 | 5.85149085 |
| Adipoq | Adiponectin, C1Q And Collagen Domain Containing | 4.594791601 | 4.63868612 |
| Sgk1 | Serine/threonine-protein kinase | 0.611314515 | 4.00199938 |
| Rbp4 | Retinol Binding Protein 4 | 1.70827297 | 3.88132035 |
| Myc | Myelocytomatosis oncogene | 0.819341704 | 3.85776255 |
| Rgcc | Regulator Of Cell Cycle | 0.557283549 | 3.68874709 |
| Orm1 | Orosomucoid 1 | 3.29037894 | 3.30270992 |
| Stra6 | Signaling Receptor And Transporter Of Retinol | 1.164271291 | 3.18095352 |
| Angptl2 | Angiopoietin-like protein 2 | 0.347129044 | 2.98379838 |
| Bmal1 | Basic Helix-Loop-Helix ARNT Like 1 | -0.831754026 | 2.90071497 |
| Cdk6 | Cyclin-dependent kinase 6 | -0.639155556 | 3.84606714 |
| Serpina3i | Cyclin-dependent kinase 6 | -4.092363499 | 3.00595597 |
| Clock | Clock circadian regulator | -0.336020299 | 3.06964803 |

**Supplementary Table S2:** Metadata for selected healthy reference (HRT) and acute kidney injury (AKI) samples from Kidney Precision Medicine Project (KPMP)

| Sample ID | KPMP ID | Disease Category | Age | Gender | Race | Procedure Type | Specimen Type | Significant Past Medical History |
|---|---|---|---|---|---|---|---|---|
| S-2008-006028 | 163-1 | HRT | N/A | N/A | N/A | Kidney Biopsy | Kidney Cortex | available on request |
| S-2008-007631 | 164-1 | HRT | N/A | N/A | N/A | Kidney Biopsy | Kidney Cortex | available on request |
| S-2108-005528 | 165-12 | HRT | 58 | Female | White | Kidney Biopsy | Kidney Cortex | available on request |
| S-2109-023337 | 165-10 | HRT | 57 | Female | White | Kidney Biopsy | Kidney Cortex | available on request |
| S-2111-008287 | 163-7 | HRT | 72 | Female | White | Kidney Biopsy | Kidney Cortex | available on request |
| S-2111-008311 | 164-12 | HRT | 37 | Female | White | Kidney Biopsy | Kidney Cortex | available on request |
| S-2112-014329 | 164-14 | HRT | 48 | Male | White | Kidney Biopsy | Kidney Cortex | available on request |
| S-2111-008287 | 164-16 | HRT | 59 | Male | White | Kidney Biopsy | Kidney Cortex | available on request |
| S-2006-001713 | 30-10868 | AKI | 55 | Male | White | Kidney Biopsy | Kidney Cortex | available on request |
| S-2006-001760 | 30-11084 | AKI | 60 | Female | White | Kidney Biopsy | Kidney Cortex | available on request |
| S-2006-001901 | 32-10456 | AKI | 41 | Male | Black or African-American | Kidney Biopsy | Kidney Cortex | available on request |

| | | | | | | | | |
|---|---|---|---|---|---|---|---|---|
| S-2006-001948 | 30-11101 | AKI | 46 | Male | Black or African-American | Kidney Biopsy | Kidney Cortex | available on request |
| S-2006-001995 | 32-10333 | AKI | 46 | Male | Asian | Kidney Biopsy | Kidney Cortex | available on request |
| S-2006-002396 | 33-10376 | AKI | N/A | N/A | N/A | Kidney Biopsy | Kidney Cortex | available on request |
| S-2006-002492 | 33-10331 | AKI | 30 | Female | African American | Kidney Biopsy | Kidney Cortex | available on request |
| S-2006-004487 | 34-10209 | AKI | 53 | Male | White | Kidney Biopsy | Kidney Cortex | available on request |
| S-2101-003289 | 30-11093 | AKI | 20 | Male | White | Kidney Biopsy | Kidney Cortex | available on request |
| S-2101-003336 | 30-11081 | AKI | 73 | Female | White | Kidney Biopsy | Kidney Cortex | available on request |
| S-2102-003410 | 34-10240 | AKI | 60 | Female | White | Kidney Biopsy | Kidney Cortex | available on request |
| S-2102-003504 | 34-10393 | AKI | 31 | Male | White | Kidney Biopsy | Kidney Cortex | available on request |
| S-2109-011411 | 32-10346 | AKI | 44 | Male | Black or African-American | Kidney Biopsy | Kidney Cortex | available on request |
| S-2109-019203 | 34-10579 | AKI | 64 | Female | Black or African-American | Kidney Biopsy | Kidney Cortex | available on request |

| | | | | | | | | |
|---|---|---|---|---|---|---|---|---|
| S-2109-019250 | 34-10666 | AKI | 48 | Male | Black or African-American | Kidney Biopsy | Kidney Cortex | available on request |
| S-2209-009415 | 34-10730 | AKI | 24 | Male | White | Kidney Biopsy | Kidney Cortex | available on request |
| S-2304-011687 | 34-10889 | AKI | 49 | Male | Other | Kidney Biopsy | Kidney Cortex | available on request |
| S-2309-028497 | 34-11066 | AKI | 40 | Male | Black or African-American | Kidney Biopsy | Kidney Cortex | available on request |
| S-2401-002293 | 933-10372 | AKI | 66 | Male | White | Kidney Biopsy | Kidney Cortex | available on request |

**Supplementary Table S3**: Pediatric chronic kidney disease (CKD) study participants characteristics (control n=36 and CKD n=72)

| Patient Characteristics | Control | CKD |
|---|---|---|
| **Age (mean)** | 11.2 | 12.6 |
| **Female gender (%)** | 50% | 45.80% |
| **Race** | | |
| *American Indian* | 0% | 0% |
| *Asian* | 2.70% | 8.45% |
| *African American* | 22.20% | 29.57% |
| *Hispanic* | 16.60% | 9.85% |
| *White* | 58% | 52.11% |
| *Hawaiian* | 0% | 0% |

**Supplementary Table S4:** End stage kidney disease study participants characteristics (n = 29).

| Age (Year) | Gender | Pre_WT (Kg) | Post-WT (Kg) | URR | Pre_BUN (mg/dL) | Post_BUN (mg/dL) | Creatinine Serum (mg/dL) | Pre_ Quinolinic acid (µM) | Post_ Quinolinic acid (µM) |
|---|---|---|---|---|---|---|---|---|---|
| 64 | M | 90.50 | 88.10 | 78.0 | 59.0 | 13.0 | 6.9 | 2.346811 | 0.03483 |
| 82 | F | 73.50 | 71.60 | 81.4 | 43.0 | 8.0 | 6.1 | 11.10638 | 1.094067 |
| 72 | F | 62.10 | 59.70 | 90.3 | 31.0 | 3.0 | 5.6 | 3.7689 | 0.023947 |
| 43 | F | 81.40 | 77.10 | 82.4 | 51.0 | 9.0 | 7.6 | 7.892067 | 0.466936 |
| 49 | M | 84.20 | 80.00 | 73.2 | 56.0 | 15.0 | 10.6 | 4.298414 | 0.567978 |
| 51 | M | 75.10 | 73.00 | 77.0 | 61.0 | 14.0 | 9.9 | 7.196925 | 0.144499 |
| 56 | F | 61.30 | 60.20 | 85.0 | 60.0 | 9.0 | 9.1 | 4.104453 | 0.02988 |
| 36 | M | 94.80 | 92.40 | 70.0 | 80.0 | 24.0 | 14.6 | 15.22459 | 4.155645 |
| 67 | F | 75.70 | 73.50 | 74.5 | 51.0 | 13.0 | 9.1 | 10.69891 | 0.470443 |
| 56 | M | 61.50 | 57.20 | 80.0 | 25.0 | 5.0 | 7.8 | 9.606912 | 0.389974 |
| 59 | F | 74.80 | 71.40 | 86.7 | 60.0 | 8.0 | 7.8 | 5.59861 | 0.022882 |
| 58 | M | 52.30 | 50.30 | 89.4 | 66.0 | 7.0 | 11.9 | 4.251141 | 0.045559 |
| 47 | F | 85.90 | 81.10 | 84.0 | 71.0 | 11.0 | 7.7 | 4.16625 | 0.130175 |
| 25 | F | 101.50 | 97.90 | 74.5 | 51.0 | 13.0 | 11.9 | 7.463583 | 0.361765 |
| 74 | M | 84.70 | 82.50 | 86.0 | 43.0 | 6.0 | 7.4 | 5.453344 | 0.197385 |
| 73 | M | 70.50 | 68.10 | 81.7 | 71.0 | 13.0 | 10.4 | 8.040016 | 0.057851 |
| 63 | F | 76.30 | 73.00 | 88.0 | 92.0 | 42.0 | 7.5 | 5.23093 | 0.136084 |
| 68 | F | 62.60 | 61.10 | 88.2 | 34.0 | 4.0 | 7.3 | 3.941061 | 0.047809 |
| 60 | F | 93.80 | 90.30 | 82.0 | 57.0 | 10.0 | 9.8 | 9.578379 | 0.451747 |
| 53 | F | 88.50 | 85.50 | 80.0 | 36.0 | 7.0 | 7.4 | 10.43136 | 0.907096 |
| 54 | F | 94.30 | 91.60 | 78.0 | 52.0 | 10.0 | 9.01 | 11.29416 | 1.023004 |
| 63 | F | 80.60 | 78.80 | 91.0 | 47.0 | 4.0 | 7.9 | 8.134879 | 0.26983 |
| 69 | F | 58.50 | 58.30 | 88.0 | 35.0 | 4.0 | 4.9 | 12.18456 | 0.288326 |
| 32 | F | 112.40 | 110.60 | 78.0 | 76.0 | 16.0 | 9.1 | 10.57339 | 0.81875 |
| 52 | F | 96.20 | 93.00 | 75.0 | 48.0 | 9.0 | 10.1 | 4.757922 | 0.204565 |
| 59 | M | 74.90 | 72.90 | 78.0 | 67.0 | 9.0 | 8.5 | 6.706208 | 0.525585 |

| | | | | | | | | | |
|---|---|---|---|---|---|---|---|---|---|
| 68 | M | 76.90 | 73.80 | 80.0 | 83.0 | 13.0 | 11.5 | 3.815608 | 0.107667 |
| 55 | M | 158.50 | 153.80 | 70.0 | 62.0 | 20.0 | 8.7 | 6.831568 | 0.69637 |
| 47 | F | 45.40 | 44.00 | 88.9 | 45.0 | 5.0 | 6.5 | 4.286252 | 0.063804 |

WT, weight; URR, urea reduction rate, BUN, blood urea nitrogen

**Supplementary Table S5:** Data analysis of reduction ratios of BUN and kynurenine pathway metabolites after a single hemodialysis session in the end stage kidney disease study

| | **URR** | **QARR** | **3HKRR** | **KYNRR** | **3HAARR** | **TRP RR** | **xant RR** | **KYNARR** |
|---|---|---|---|---|---|---|---|---|
| **Number of values** | 29 | 29 | 28 | 28 | 28 | 28 | 28 | 28 |
| **Minimum** | 70.00 | 72.70 | 18.56 | 9.788 | 48.59 | -104.1 | 23.72 | 10.56 |
| **Maximum** | 91.00 | 99.59 | 97.85 | 94.52 | 91.44 | 68.28 | 70.33 | 72.15 |
| **Range** | 21.00 | 26.89 | 79.29 | 84.73 | 42.85 | 172.4 | 46.61 | 61.60 |
| **Mean** | 81.35 | 94.84 | 85.10 | 56.61 | 71.27 | -5.691 | 46.28 | 43.37 |
| **Std. Deviation** | 6.064 | 5.440 | 16.46 | 16.21 | 9.837 | 37.05 | 13.48 | 15.00 |
| **Std. Error of Mean** | 1.126 | 1.010 | 3.111 | 3.064 | 1.859 | 7.001 | 2.548 | 2.834 |
| **P-value (two tailed t test)** | 5.1913E-19 | 1.22487E-15 | 3E-12 | 2E-09 | 1.6E-11 | 0.51887 | 6.8E-09 | 0.002115748 |

Reduction ratio (RR) represents the percent decrease in each analyte post-dialysis. QA, quinolinic acid; 3HK, 3-hydroxykynurenine; KYN, kynurenine; 3HAA, 3-hydroxyanthranilic acid; Trp, tryptophan; Xant, xanthurenic acid; KYNA, kynurenic acid; BUN, blood urea nitrogen.

### Supplementary Methods

**RNA sequencing.** RNA quality was assessed using the Agilent Tapestation 4200, and only samples with an RNA Integrity Number (RIN) ≥ 8 were included for sequencing. RNA concentration was measured with a Qubit® 4.0 Fluorimeter (ThermoFisher). One microgram of DNase-treated total RNA was used for library preparation with the TruSeq Stranded mRNA Library Prep Kit (Illumina). Poly(A) RNA was purified and fragmented, followed by strand-specific cDNA synthesis. The resulting cDNA was A-tailed, ligated to indexed adapters. After adapter ligation, samples are PCR amplified and purified with AmpureXP beads, then validated again on the Agilent Tapestation 4200. Before being normalized and pooled, samples are quantified by Qubit then run on the Illumina NextSeq 2000 on a P2-100 flowcell. Raw sequencing data were processed with bcl2fastq to generate FASTQ files. Gene-level read counts were obtained using featureCounts, and differential expression analysis was performed with edgeR, using the mm10 reference genome.

**Supplementary Figure S2.** Eight-week-old WT C57BL/6J mice were used for this study. All experimental procedures were approved by the Institutional Animal Care and Use Committee at the University of Texas Southwestern Medical Center and conducted in accordance with the NIH Guide for the Care and Use of Laboratory Animals. Animals were randomly assigned into three groups. Control group (n = 5): Mice received standard drinking water for 14 days and were injected intraperitoneally with folic acid (FA, 250 mg/kg) on day 0. Animals were sacrificed on day 14. NAM-treated group (n = 8): Mice were administered 0.6% nicotinamide (NAM) in drinking water starting on day 0 and were injected intraperitoneally with FA (250 mg/kg) on day 0. Drinking water was exchanged every other day. Animals were sacrificed on day 14.

**qPCR primers list**

Mouse_*Qprt:*

F_ GAAAGACAACCATGTAGTGGCGG; R_ GGCTGCTACATTCCACCTCTAC

Mouse_*Col3a1:*

F_GACCAAAAGGTGATGCTGGACAG; R_CAAGACCTCGTGCTCCAGTTAG

Mouse_*Tgfb*:

F_TGATACGCCTGAGTGGCTGTCT; R_CACAAGAGCAGTGAGCGCTGAA

Mouse_*Acta2:*

F_TGCTGACAGAGGCACCACTGAA; R_CAGTTGTACGTCCAGAGGCATAG

Mouse_*Lcn2:*

F_ ATGTCACCTCCATCCTGGTCAG; R_GCCACTTGCACATTGTAGCTCTG

Mouse_*Nfkb:*

F_ CCCCCTGGTGGAGAACTTTG; R_ GCTGCTTCATGTCCCCTTGT

Mouse_*Pdgfra:*

F_ CATGCGGGTGGACTCTGATA; R_ CCGAAGTCTGTGAGCTGTGT

Mouse_*Gapdh:*

F_TGTGTCCGTCGTGGATCTGA; R_CCTGCTTCACCACCTTCTTGAT

Mouse_*Actin:*

F_CATTGCTGACAGGATGCAGAAGG; R_TGCTGGAAGGTGGACAGTGAGG
